# ChlORIS: Chloroplast Orthologs Resource & Identification Suite

**DOI:** 10.64898/2026.08.05.743164

**Authors:** Yuhao Tong, Vanessa Rossetto Marcelino, Robert Turnbull, Heroen Verbruggen

## Abstract

Chloroplast or plastid genomes are essential resources for studying the evolution and diversity of algae and land plants. Although thousands of plastid genomes have been sequenced, their full potential has not been realised; derived resources such as orthogroup databases and reference datasets for metagenomic profiling remain underdeveloped. We present the ChlORIS database to address these problems across all algal phyla. From 2,254 publicly available algal plastid genomes, after dereplication we clustered 2,531 orthogroups from the annotated proteins and selected 496 orthogroups with consistent gene naming, enabling cross-genome comparisons of homologous plastid proteins. We further selected 224 core orthogroups, each containing more than 10 protein sequences, for which we produced score-calibrated hidden Markov models (HMMs), multiple sequence alignments and predicted protein structures. The value of these resources for phylogenomics is demonstrated through a large-scale plastid phylogeny of 859 taxa spanning all major algal lineages. We characterised the protein HMMs by cross-referencing them to Pfam domains and calibrated score cutoffs for reliable detection. The metagenomic database, HMM library, nucleotide and amino acid alignments, predicted structures and protein metadata, cross-linked to UniProt and InterPro (Pfam), are openly available on the ChlORIS website at https://chloris.codeberg.page/.

## Introduction

Algae are critical to the functioning of aquatic ecosystems. Their photosynthesis accounts for nearly half of global net primary production and oxygen generation, and contributes substantially to oceanic carbon uptake (Field et al. 1998). Multicellular algae (seaweeds) are foundation species that provide habitat for coastal marine species (Wernberg et al. 2011) and underpin coastal ecosystem goods and services valued at hundreds of billions of dollars annually (Eger et al. 2023). These photosynthetic eukaryotes span diverse morphologies, ranging from unicellular protists, over small filamentous or colonial forms, to large macroscopic organisms like kelps and seaweeds (Guiry 2012). Their emergence dates back approximately 1.5 billion years (Yoon et al. 2004). There are roughly fifty thousand formally described species of algae today, spanning 14 phyla across several kingdoms including Plantae, Chromista and Protozoa (Guiry 2024), and considerable additional diversity remains cryptic or undescribed, with environmental sequencing continually uncovering novel lineages. A key resource to study their evolution and physiology is the plastid genome: a reduced derivative of the endosymbiotic cyanobacterial genome that encodes essential photosynthetic protein machinery along with some other functions (De Vries and Archibald 2018). The genome structure and protein-coding genes are generally conserved, making them well-suited for profiling evolutionary relationships and sampling novel algal taxa (Jackson et al. 2018).

The number of available plastid genomes is growing rapidly, with many studies using newly sequenced plastid genomes to derive insights into algal evolution and plastid genome structure (Kamikawa et al. 2015; Füssy et al. 2025). Many such studies also re-use previously sequenced genomes, but do so relying on *ad hoc* pipelines, posing reproducibility challenges. Databases such as OGDA (Organelle Genome Database for Algae; Liu et al. 2020) have been developed for sequenced organellar genomes, storing metadata with direct links to the genomes on NCBI. However, these resources do not summarize information on protein families or the evolutionary relationships of individual genes, limiting their utility for certain applications. Meanwhile, global microbial profiling efforts (e.g., MicrobeAtlas; Matias Rodrigues et al. 2026) have revealed numerous novel plastid sequences of microeukaryotes in environmental sequencing data, but detecting and analysing these is problematic due to a lack of plastid-specific reference datasets for use in metagenomic workflows for evaluating metagenome-assembled genomes (MAGs), such as CheckM (Parks et al. 2015).

To address these gaps, we present the ChlORIS database: a unified data resource for plastid orthologs. ChlORIS supports multiple aspects of plastid genomics and evolutionary research, including orthogroup clusters and multiple sequence alignments for phylogenetics and molecular evolution. We also provide profile hidden Markov models (HMMs) for remote homology detection in metagenomic and other novel datasets, and a CheckM-compatible database for MAG quality assessment. Compared with previous plastid genomic resources, ChlORIS is the most comprehensive resource, integrating orthogroup classifications, protein domain annotations, and quality-scored HMMs, with direct applications to evolutionary biology, metagenomics, and biotechnology. All resources are openly available and cross-linked to UniProt and InterPro (Pfam).

## Methods

The database construction workflow is separated into three sections: data curation, data analyses, and metagenomic database construction (Fig. 1).

**Figure 1.**
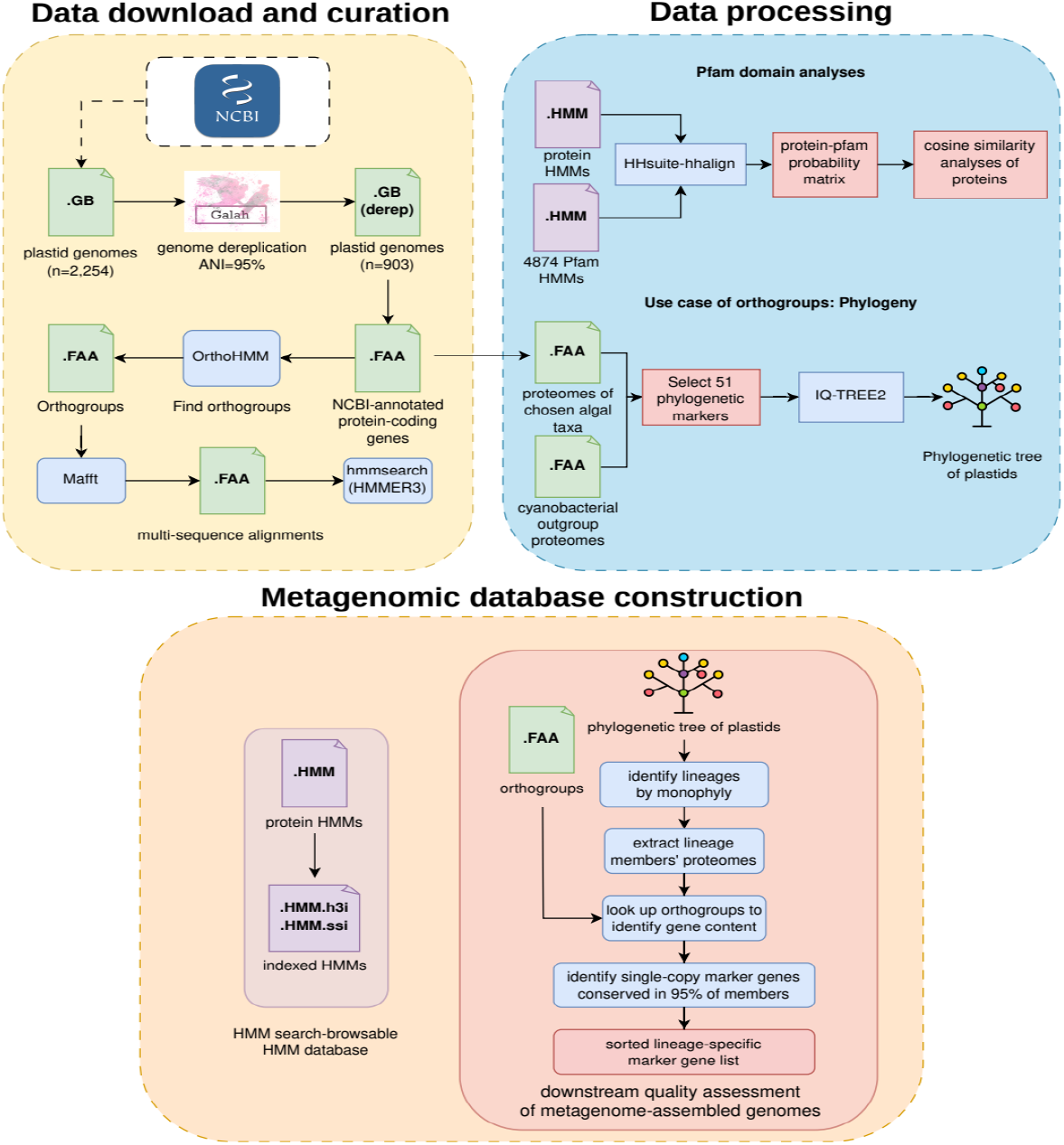
ChlORIS database construction workflow. First, we prepared the source data by downloading them from NCBI GenBank, filtering only protistan algae and macroalgae plastids while removing derived plant plastids and kleptoplastic plastids in sacoglossan sea slugs. We dereplicated the genomes, extracted the proteomes from them and clustered orthogroups via OrthoHMM. Alignment files and orthogroup profile HMMs were derived. We then analysed protein domain content of selected HMMs and performed a phylogenetic analysis to demonstrate the utility of the orthogroups. We also reformatted our HMMs and used the generated phylogeny to construct a metagenomic database in a format compatible with CheckM, enabling plastid detection in environmental sequencing data.

### Genome preparation

We downloaded from 2,254 plastid genomes of micro- and macroalgae from GenBank. We excluded land plants, other derived plant plastids, and kleptoplasts of sacoglossan sea slugs. The genomes were dereplicated at 95% average nucleotide identity (ANI) using Galah v.0.4.2 (Aroney et al. 2024), reducing the total number of genomes to 903. We discarded genomes without annotations, retaining 893 genomes.

### Orthogroup inference and alignments

We extracted genes with annotated amino acid sequences from each retained genome. Then, we used OrthoHMM v.0.1.0 (Steenwyk et al. 2024) with default parameters to cluster the extracted proteins into orthogroups: groups of homologous plastid-encoded gene sequences. Through manual curation, we identified orthogroups with consistent annotation names, excluding those dominated by hypothetical open reading frames (ORFs), which typically lack consistent naming. We produced protein (amino acid) alignments of all orthogroups using the localpair algorithm in MAFFT v.7.526. CDS extracted from source GenBank records were back-translated onto the existing columns of the protein alignments, producing codon-level alignments maintaining the same gaps as the protein alignment, though as triplets.

### Curation of HMMs

To assign quality scores to each protein HMM, we calibrated them against the orthogroup assignments. Using orthogroup assignments as ground truth, we determined true positives, false positives, and false negatives for each HMM. Next, we ran hmmsearch (HMMER v.3.4) on the proteome using each named gene’s HMM (Eddy 2011). We computed noise cutoffs (NC) and trusted cutoffs (TC), score thresholds that distinguish reliable hits from background noise, based on the bit scores of the highest-scoring false positive and lowest-scoring true positive, respectively. These scores are then added to each protein HMM.

### Protein structure and Pfam domains

To annotate each protein with its Pfam domains and to characterise domain-sharing among plastid proteins, we linked our full-length protein HMMs to the Pfam database r27 (Mistry et al. 2021). We performed HMM–HMM alignments using HHalign (Steinegger et al. 2019) between 496 protein HMMs and 4,874 Pfam HMMs. We converted both the protein orthogroup and Pfam domain data to the HH-suite-format to perform HMM-HMM comparison, aligning them with hhalign (HH-suite v.3.3.0). We derived probability tables recording the likelihood of each Pfam domain occurring in each protein. We then quantified pairwise protein similarity based on their shared Pfam domain profiles using cosine similarity, retaining proteins and Pfam domains with significant hits. Using these data, we plotted the protein cosine similarity heatmaps for 9 plastid functional groups: photosystem complex I (*psa*), photosystem complex II (*psb*), ATP synthase complex (*atp*), cytochrome b6f complex (*pet*), ribosomal protein small subunits (*rps*), ribosomal protein large subunits (*rpl*), RNA polymerase (*rpo*), phycobilisome subunits (*apc, cpc* and *cpe*) and conserved hypothetical proteins (*ycf*). For each selected orthogroup we predicted a representative protein structure with AlphaFold 3 (Abramson et al. 2024) from the orthogroup alignment, and provide the models in CIF format on the website.

### Calibrating HMM score cutoffs

We characterised how each HMM recovers the members of its own orthogroup, using the OrthoHMM clustering as the reference. The ca. 116,000 clustered plastid proteins (section 2.2) formed the reference set, and for the 224 orthogroups with more than 10 members we ran hmmsearch against it at an e-value cutoff of 10^-3^, labelling each hit as a correct or off-target assignment relative to the clustering. For each HMM we recorded precision (the proportion of hits belonging to the target orthogroup), recall (the proportion of orthogroup members detected) and their harmonic mean, the F1 score, which we used to flag orthogroups prone to cross-hits between paralogs, and we set the noise (NC) and trusted (TC) cutoffs as described in section 2.3.

### Phylogenetics

We conducted a phylogenetic analysis to demonstrate the utility of the orthogroup database and evaluate phylogenetic signal. To this goal, we downloaded 7 cyanobacterial genomes including the *Gloeomargarita* lineage considered the closest extant relative of the plastid donor for the principal primary endosymbiosis event (Ponce-Toledo et al. 2017). For the tree (but not the ChlORIS database) we pruned 34 non-photosynthetic genomes, since their lower gene content and accelerated rates of sequence evolution may destabilise their placement. The remaining 859 photosynthetic plastid genomes, together with 7 cyanobacterial outgroups (866 taxa in total), were used to build a supermatrix of 51 conserved marker genes. We first aligned individual protein sequences using the localpair algorithm in MAFFT v.7.526 (Katoh 2002) and trimmed them using ClipKIT v.2.7.0 (Steenwyk et al. 2020). We then concatenated the single-protein alignments with PhyKIT v.2.1.2 (Steenwyk et al. 2021) and ran IQTREE v2.3.6 (Nguyen et al. 2015) with 1000 ultrafast bootstrap replicates and automated model selection (MFP+MERGE).

### Quality-assessment database for plastid MAGs

We further extend ChlORIS by formatting it as a database compatible with CheckM for assessing the quality of metagenome-assembled genomes, including completeness (the proportion of expected genome content present in the MAG) and contamination (the proportion of sequences that may originate from other organisms). This uses the HMMs formatted for use with CheckM along with a gene content database across taxonomic lineages. Using the phylogenetic tree from section 2.6, we identified 39 branches (hereafter lineages) with monophyly after mapping leaves to their taxonomy. For each lineage, we extracted lineage members’ proteomes and mapped genes to orthogroups to determine gene content. By applying the criteria from CheckM (Parks et al. 2015) with modifications, we selected marker genes present as single copy in >95% of members and grouped them into marker gene sets used to infer completeness and contamination.

## Results

### ChlORIS database contents

ChlORIS version 1.0 contains 2,531 orthogroups from 893 input plastid genomes. Through manual curation, we selected 496 orthogroups for which most members shared a standard gene name (e.g., *psa*A, *atp*B, *rpl*4). We refer to this 496-orthogroup set as the “full” database. The remaining 2,035 orthogroups, including 1,229 singletons, are mostly uncharacterised hypothetical proteins without consistent gene names. We also defined a “core” set of 224 genes, which is a subset of the 496 orthogroups containing >10 aligned sequences. This set corresponds to the major proteins conserved across all or large algal phyla.

Aligning 4,874 hidden Markov models from Pfam-A against the HMMs of our 496 orthogroup set resulted in most orthogroups (470) returning at least one significant Pfam hit (involving 989 Pfam domains in total), providing homogeneous and reproducible functional annotation of the orthogroups independent of the original GenBank entries.

### Calibrated cutoffs reduce off-target hits

For the core gene set, we examined how each of the 224 protein HMMs recovered its own orthogroup members among the total pool of plastid proteins analysed here (ca. 116,000), using the original orthogroup assignments as ground truth. Using a simple e-value cutoff (10^-3^), most HMMs recovered their members cleanly, yet there were some outliers with F1 scores ≤0.8, including some off-target hits (Fig. 2a,b). This reflects expected relationships among functionally related plastid proteins that have domain-sharing among paralogous proteins. We illustrate this for the documented cases of gene families that have diversified through duplication events, including the phycobiliprotein superfamily (Fig. 2a) (Apt, Collier, and Grossman 1995) and the D1/D2 (*psb*A/*psb*D) and CP47/CP43 (*psb*B/*psb*C) pairs of photosystem II (Fig. 2b) (Cardona 2015). Applying calibrated noise and trusted cutoffs removed most of these off-target hits, leaving only a few proteins with F1 score ≤0.85 (Fig. 2c).

**Figure 2.**
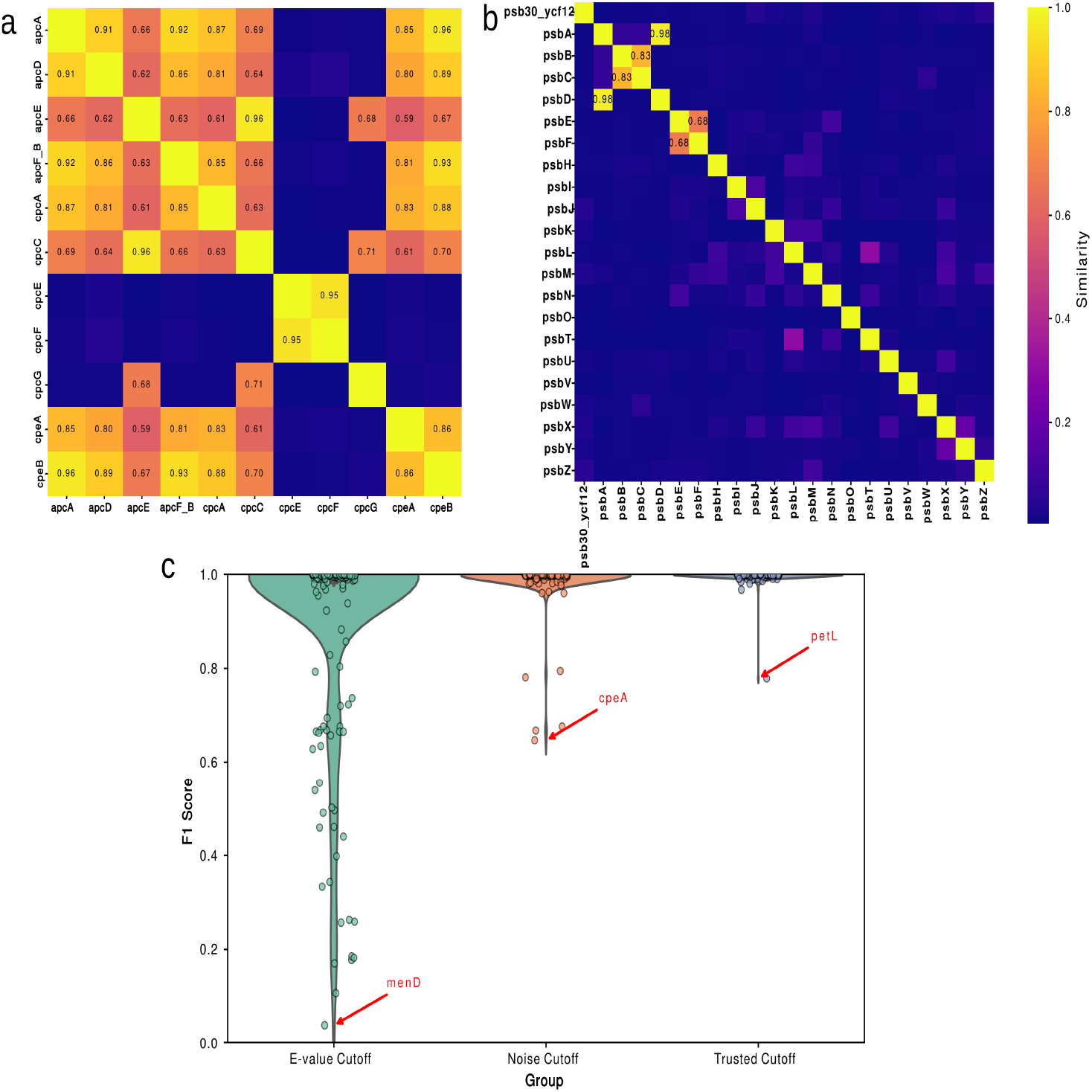
Protein HMM domain similarity and off-target hits using e-value of 10^-3^ and noise cutoff. (a & b) Protein-to-protein domain similarity measured by the cosine similarity for each pair of proteins in phycobilisome complex (a) and photosystem II (b), with the probability calculated by HH-suite’s hhalign function. (c) The distribution of F1 score of protein HMMs using three cutoffs: e-value at 10^-3^, noise cutoff and trusted cutoff when running hmmsearch.

### Phylogeny of 859 plastid genomes

As an example of using these orthogroups, we constructed a large-scale phylogenetic tree from 51 conserved genes obtained from 859 plastid genomes from the database (non-photosynthetic species were excluded) and 7 cyanobacterial outgroups, spanning nearly all major primary and secondary plastid lineages across the eukaryotic tree (Fig. 3). Our orthogroups recover the expected evolutionary signal across the plastid genome phylogeny, with high concordance to the placement of phyla in previous studies (Jackson et al. 2018; Figueroa-Martinez et al. 2019; Kim et al. 2017; Yang et al. 2016).

**Figure 3.**
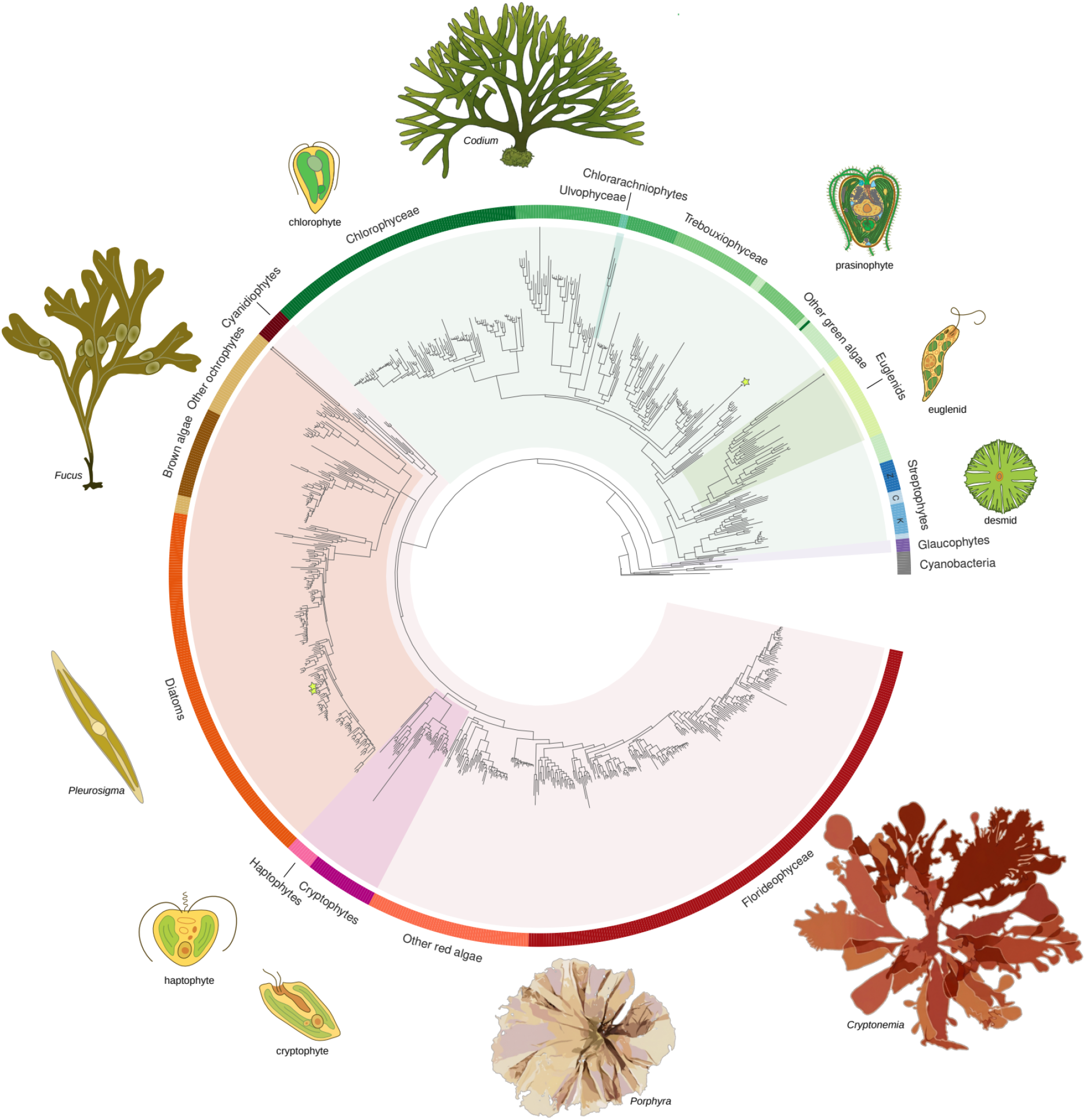
Plastid phylogenomic tree of 859 eukaryotic algae and 7 cyanobacterial outgroups. Background tints indicate primary plastid lineages (red/green/purple) with nested secondary endosymbiotic clades. Small open circles at the end of a branch indicate the branch was truncated for visualisation. Z = Zygnematophyceae + land plants, K = Klebsormidiophyceae, C = other charophytes. Green stars indicate tertiary or serial secondary dinoflagellate endosymbioses. Image credits: Original illustrations are by Patrick Keeling, Yana Eglit (chlorophyte, prasinophyte, euglenid, desmid, haptophyte, cryptophyte; (Keeling and Eglit 2023)) and Heroen Verbruggen (Fucus and Pleurosigma, via Figshare; from (Bringloe et al. 2020)). The images of Cryptonemia, Porphyra and Codium are LLM-rendered representations based on public domain photographs from the Auckland Museum (Wendy Nelson, Cryptonemia), Luis Fernández García (Porphyra), Cwmhiraeth (Codium), all accessed via WikiMedia Commons.

### Data availability and the ChlORIS website

We provide ChlORIS as a resource that is fully in the public domain, under a Creative Commons Zero (CC0) license permitting any type of use. The predicted protein structures are the sole exception, since they are subject to the AlphaFold 3 terms of use.

The ChlORIS website (https://chloris.codeberg.page/, Fig. 4) is the main access point for accessing and browsing the data. The database’s scope and potential use cases are described on the “Description” page, along with brief biosketches of its developers. The genomes and genome clusters we included in these analyses are listed in a table on the “Species” page. Metadata, Pfam domain information, browsable protein alignments, and predicted structures for the 224 core genes, are given on the “Proteins” page, with an index page listing all proteins. Each protein links to a representative UniProt entry (the representative sequence of its UniRef90 cluster) and to the Pfam domain families it contains, which resolve to InterPro. A “Download” page provides bulk access to all resources, profile HMM library, multiple sequence alignments (nucleotide and protein level), predicted protein structures (CIF format) for the selected orthogroups, the unselected orthogroups, and the phylogeny (newick), and the 2,035 uncharacterised ORF orthogroups. A “Recipes” page links to defined workflows for using ChlORIS in particular applications (cf. the use cases described below).

**Figure 4.**
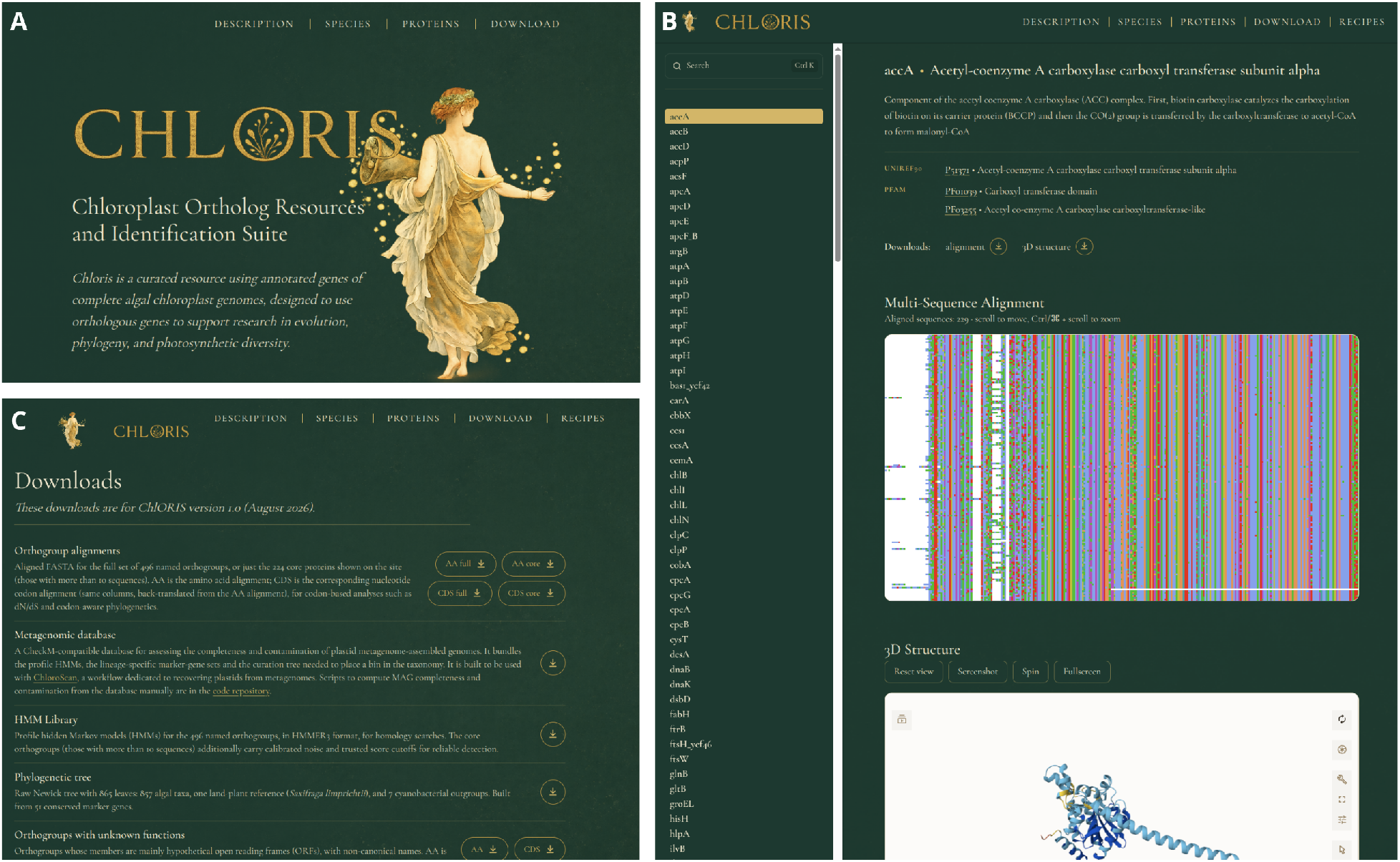
ChlORIS website screenshots. A. Landing page. B. Proteins page. C. Downloads page.

The website is hosted on CodeBerg, a non-profit forge, aligned with ChlORIS’s open-data mission. The key datasets are hosted on Zenodo (https://doi.org/10.5281/zenodo.20845450), where versioned accessions of the database will be kept for reproducibility as future updates to ChlORIS are made. The accompanying code for ChlORIS, including aspects of the workflow used for its creation, and the graft scripts (see below), are hosted on CodeBerg (https://codeberg.org/chloris) using the MIT License.

### Use case examples and recipes

We now describe two “recipes” for how ChlORIS could be used. First, in phylogenetics, the protein HMMs can be directly used to extract marker genes from new input genomes. By directly feeding HMMs and input genomes to OrthoFisher v.1.1.2 (Steenwyk and Rokas 2021), the results identify genes with significant hits. These can then be aligned, concatenated, and used for tree inference. ChlORIS itself can serve as a valuable backbone dataset for such applications, with its alignments and the our 859-tip tree serving as the backbone for the researcher’s new data to be inserted. To facilitate this kind of application, we provide the “graft” toolkit, a set of python scripts, as a solution to graft new genomes into the ChlORIS backbone in a reproducible fashion (https://codeberg.org/chloris/graft). This toolkit automates the download of relevant ChlORIS datasets, filtering by ingroup and outgroup taxa, assignment of gene sequences from a user’s new genomes to the orthogroups (hmmscan), and the placement of these sequences in the ChlORIS curated alignments without altering the existing alignment columns (mafft).The resulting alignments can then be fed into downstream phylogenomic workflows, including the option to graft the new genome into ChlORIS’s reference genome without re-inferring the backbone topology.

Second, in metagenomics, one can apply the ChlORIS database in quality control of MAGs, to determine which recovered genome bins meet the researcher’s quality thresholds for inclusion in downstream analyses. With our formatted metagenomic database, tools like CheckM (D. H. Parks et al. 2015) and binny (Hickl et al. 2022) can use the protein HMMs from ChlORIS to scan their presence and estimate the completeness and contamination levels of the MAGs. This use case is described in more detail in the description of the ChloroScan software, which employs this method (Tong et al. 2026).

## Discussion

ChlORIS brings plastid orthogroups, alignments at the protein and nucleotide (codon) level, quality-scored HMMs, predicted structures and a phylogenomic tree together in a single openly licensed resource cross-linked to UniProt and InterPro (Pfam). Beyond the two use cases above, ChlORIS supports several further applications. The quality-scored HMMs enable homology-based annotation of newly sequenced or divergent plastid genomes, and screening of eukaryotic genome assemblies for plastid contigs. The backbone phylogeny provides a reference for phylogenetic placement and taxonomic assignment of new genomes and recovered plastid MAGs. The orthogroup CDS and protein alignments, structures and tree are ready as inputs for molecular evolutionary analyses including selection, substitution rate and divergence time studies, while information about protein structure and conserved residues can feed into applied research and chloroplast engineering. The alignments also support primer design for DNA (meta-)barcoding and reference-based profiling of algal reads in environmental data. Lastly, our curated dataset offers a benchmark for orthology inference and downstream applications in phylogenomics.

Current limitations of the database include that reference genomes are mainly from lineages with canonical plastid structure. Some algae with non-canonical plastids from dinoflagellates (He et al. 2024; Howe, Nisbet, and Barbrook 2008) and Cladophorales green algae (Bjornson et al. 2026) have not yet been included in ChlORIS. Data for such lineages are scarce and their deviant nature makes them harder to fit into canonical workflows, however, we plan to include these lineages in future releases.

Dereplication at 95% ANI curbs redundancy from densely sequenced taxa but will collapse related species and therefore under-represent within-genus diversity. Furthermore, ChlORIS uses taxonomic annotations as well as gene and intron boundaries directly from source GenBank records without independent re-annotation; consequently, pre-existing errors in those records propagate into the current version of the database. A systematic re-annotation of source sequences, facilitated by the orthogroup framework built here, is planned for a future version of ChlORIS.

Two developments position ChlORIS to grow. Reference plastid genomes continue to accumulate, driven bottom-up by evolutionary and taxonomic studies (e.g., Starko et al. 2021; DíWaz-Tapia et al. 2017; Cremen et al. 2019; Hughey et al. 2026) and increasingly by biodiversity genomics initiatives such as the Darwin Tree of Life project, ATLASea, the European Reference Genome Atlas and the National Biodiversity DNA Library (CSIRO 2026; Darwin Tree of Life Consortium 2022; ATLASea 2026; Mc Cartney et al. 2024). At the same time, environmental sequencing has reshaped our understanding of microbial diversity: metagenome-derived genomes have dramatically expanded the known tree of life (Hug et al. 2016; Parks et al. 2017; Spang et al. 2015) and prokaryote genome databases such as GTDB (Parks et al. 2026) and ProGenomes4 (Fullam et al. 2026). Yet the eukaryotic, and especially organellar, fraction of environmental sequencing data remains much less explored. Plastid-targeting workflows such as ChloroScan now make this fraction accessible (Tong et al. 2026), and early studies confirm that mining novel algal plastid genomes from metagenomic data is achievable (Jamy et al. 2025; Shrestha et al. 2025), so we expect plastid MAGs to accumulate rapidly.

In the light of these new data sources, we anticipate updating ChlORIS on a yearly cycle to incorporate newly available genomes and high-quality MAGs. This will continue to increase taxonomic coverage and HMM sensitivity, further strengthening ChlORIS as a resource for metagenomics, evolutionary biology, bioengineering, and a range of other applications.

## Abbreviations

ANI: average nucleotide identity
CDS: coding sequence
CIF: Crystallographic Information File
HMM: hidden Markov model
MAG: metagenome-assembled genome
NC: noise cutoff
ORF: open reading frame
TC: trusted cutoff.

## Funding

Australian Biological Resources Study (4-G046WSD), Norwegian Taxonomy Initiative (Artsdatabanken, projects 6-24 and 86-25), Spanish Ministry of Science and Innovation (RYC2023-042907-I).

## Acknowledgements

We are grateful to the Verbruggen lab members for fruitful discussions and encouragement as we were developing this resource. We appreciate the help of Bethany Turnbull for the design of the database website. We thank Franz Goecke and Sophie Steinhagen, the lead PIs of two Norwegian Taxonomy Initiative projects, for facilitating HV’s participation in their projects, which helped support this work.

## Author contributions

Y.T.: methodology, software, formal analysis, writing – original draft; V.R.M.: methodology, supervision, writing – review & editing; R.T.: methodology, supervision, writing – review & editing; H.V.: conceptualization, supervision, software, formal analysis, writing – original draft, writing – review & editing, funding acquisition.

## Notes

https://chloris.codeberg.page

### Competing Interest Statement

The authors have declared no competing interest.

